# Chemically Synthesized Ultra-long DNA as Building Blocks to Accelerate Complex Gene Construction in Synthetic Biology

**DOI:** 10.1101/2025.04.02.646740

**Authors:** Mancang Zhang, Yang Hu, Hao Huang, Yongyong Shi

**Author notes:** These authors contributed equally to this work (Mancang Zhang, Yang Hu).

## Abstract

Certain applications of synthetic biology rely on the construction of large and complex DNA sequences. Current DNA synthesis technologies are limited in their capacity to generate ultra-long oligonucleotide for complex gene construction with extensive repetitive motifs and uneven base distribution efficiently. Here, we report a novel platform named UCOS (short for **U**ltralong **C**omplex **O**ligonucleotides **S**ynthesis) that enables the efficient synthesis of long, complex, and challenging DNA fragments. This platform employs nonporous silica microspheres as the solid support instead of traditional CPG (Controlled Pore Glass) solid support, full-length enrichment based on 5’ flank sequence hybridization and an error-removing enzyme for correct sequence selection, substantially enhancing the fidelity of intricate, ultralong oligonucleotides. Using this approach, we successfully synthesized challenging sequences up to 600nt in length, encompassing tandem repeats and uneven base distributions. Overall, this novel platform demonstrates exceptional efficiency and reliability in handling ultralong DNA fragments with highly repetitive and complex features. It provides a strong foundation for advancing synthetic biology and shows great potential as a powerful tool for constructing challenging genes and enabling the customized synthesis of functional genetic elements.

## Background

DNA, the fundamental molecule of life and carrier of genetic information, has been at the forefront of biotechnological advancement. The development of DNA synthesis technologies has revolutionized various fields, including synthetic biology^1^, drug development^2^, and gene therapy^3^. Since the inception of oligonucleotide chemical synthesis research in the 1950s^4^, significant progress has been made. The phosphodiester method was proposed by H. G. Khorana and was a common method for the early synthesis of oligonucleotides^5^. A notable breakthrough came in the 1980s when Marvin Caruthers reported the phosphoramidite method for oligonucleotide synthesis^6^. Currently, the solid-phase phosphoramidite method, developed by Beaucage and Caruthers, remains the most widely used approach, comprising four essential steps: deprotection, coupling, capping, and oxidation^7^.

The evolution of oligonucleotide synthesis capabilities has opened new frontiers in molecular biology^8, 9^. The ability to synthesize 20-30 base sequences enabled the development of PCR and DNA sequencing, establishing foundations for recombinant DNA technology and molecular diagnostics. Furthermore, the capacity to synthesize 50-100 base sequences facilitated precise DNA manipulation techniques, including site-directed mutagenesis and genetic engineering.

However, chemical DNA synthesis has reached a plateau due to inherent limitations^10^. Current synthesis processes face challenges such as depurination^11^ caused by chemical reagents and incomplete capping, making the synthesis of oligonucleotides longer than 200 nucleotides particularly challenging^8, 12^. Longer DNA sequences are typically assembled through the “splicing” of shorter oligonucleotides. This process involves synthesizing fragments less than 100 nucleotides (typically 80nt) with homologous ends, followed by assembly using techniques such as PCA^13, 14^ or Gibson assembly^15-17^. Nevertheless, this approach encounters difficulties with challenging sequences, particularly those containing long repetitive elements or unbalanced base composition.

The limitations of chemical synthesis have prompted investigation into enzymatic DNA synthesis technologies, building upon the discovery of DNA polymerases and advances in synthetic biology. Terminal deoxynucleotidyl transferase (TdT)^18, 19^ has emerged as a promising template-independent polymerase for de novo DNA synthesis. However, enzymatic synthesis faces its own challenges, including high costs associated with enzyme-nucleotide conjugate preparation and the requirement for modified nucleotides. Additionally, the base composition of synthesized sequence termini, particularly the last two bases, significantly influences nucleotide incorporation efficiency through base stacking interactions. Studies^20^ have shown poor incorporation efficiency of 3′-ONH_2_-dATP by TdT, especially with CT terminal bases, highlighting sequence-dependent limitations of enzymatic synthesis.

Building upon these challenges in DNA synthesis, particularly for sequences with unique structural features, there is an urgent need for innovative solutions. The complexity of target sequences in modern biotechnology applications presents diverse synthesis challenges that require novel approaches.

Among the most demanding targets are sequences with extreme base composition bias and repetitive elements. For example, the Plasmodium falciparum genome, with its unusually high AT content (approximately 80-85%)^21^, represents a fundamental challenge in synthesis fidelity and assembly. Similarly, spider silk protein genes, characterized by their extensive repetitive modules and substantial length (often exceeding 10kb), contain multiple iterations of poly-alanine blocks and glycine-rich regions that are crucial for biomaterial engineering applications^22^. The synthesis of Tandem Repeat (TR) sequences^23^, essential for genetic fingerprinting and disease research, presents additional complexity due to their highly repetitive nature. Further challenges include G-quadruplex forming sequences in telomeres^24^, large structural RNAs^25^, and synthetic genes containing complex secondary structures^26^.

To address these specific challenges, we have developed a novel synthesis platform optimized for Ultra-long Complex Oligonucleotides Synthesis (UCOS). A schematic representation of the synthetic process is shown in Figure 1. Our approach innovatively combines the use of nonporous solid silica microspheres as synthesis carriers (Figure 1A) instead of CPG, which significantly enhance coupling efficiency and stability for ultralong oligonucleotide; Furthermore, we use biotinylated primers to enrich full-length fragments and filter out incomplete products (Figure 1B). Additionally, mismatch repair enzymes are employed to remove fragments containing synthesis errors, such as insertions, deletions, and base substitutions (Figure 1C). The high-accuracy full-length oligonucleotides obtained are then further assembled into vectors for subsequent functional validation (Figure 1D).

**Figure 1.**
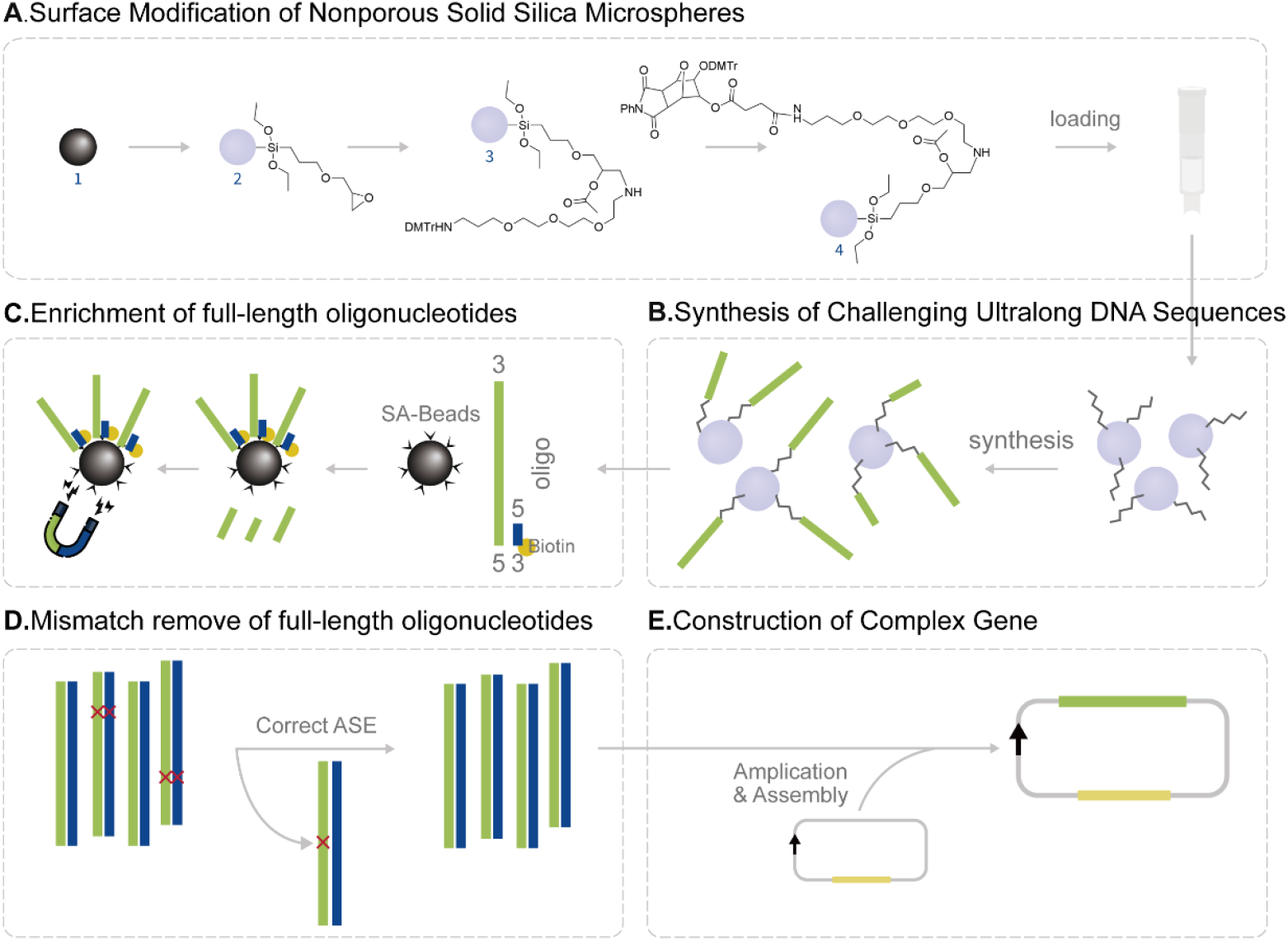
The workflow of Ultra-long Complex Oligonucleotides Synthesis (UCOS) Platform. (A). Schematic illustration of the surface modification of Nonporous Solid Silica Microspheres. Primary surface grafting of pretreated raw nonporous solid silica microspheres to yield intermediate 2; Secondary grafting and capping of residual hydroxyl groups to prevent unwanted downstream reactions to produce intermediate 3; Deprotection of DMT groups followed by condensation reaction with linker molecules to generate product 4; Column packing with product 4 to prepare the solid support for subsequent oligonucleotide synthesis. (B). Schematic representation of ultra-long DNA fragment synthesis occurring in the packed synthesis column. (C). Selection of full-length Oligonucleotides. To screen the full-length synthetic oligonucleotides, we designed and synthesized a biotinylated primer complementary to the 5’ termini of the target sequences. Following annealing with the biotinylated primers, full-length sequences were captured using streptavidin-coated magnetic beads. (D). Mismatch remove of the enriched full-length Oligonucleotides. mismatch repair enzymes were used to eliminate the error-containing sequences. (E). Schematic representation of vector construction. Following PCR amplification, the sequences were cloned into plasmid vectors for validation studies.

Using this platform, we have successfully synthesized challenging sequences up to 600 nucleotides in length, including those with repetitive elements and unbalanced base composition. The biological functionality and sequence accuracy of these synthesized oligonucleotides have been thoroughly validated. This advancement represents a significant step forward in enabling applications across cellular and gene therapy, protein engineering, biomanufacturing, and fundamental life science research.

## Results

### Design and Efficient Synthesis of Challenging DNA Sequences

To validate our established platform for synthesizing challenging ultra-long sequences, we initially attempted to synthesize poly(T) sequences of varying lengths. Multiple synthesis reactions were designed to produce fragments ranging from 200 to 400 nucleotides (nt). Polyacrylamide gel electrophoresis (PAGE) analysis of the purified products revealed single bands of expected sizes for all samples (Figure 2A), confirming the successful synthesis of the designed sequences.

**Figure 2.**
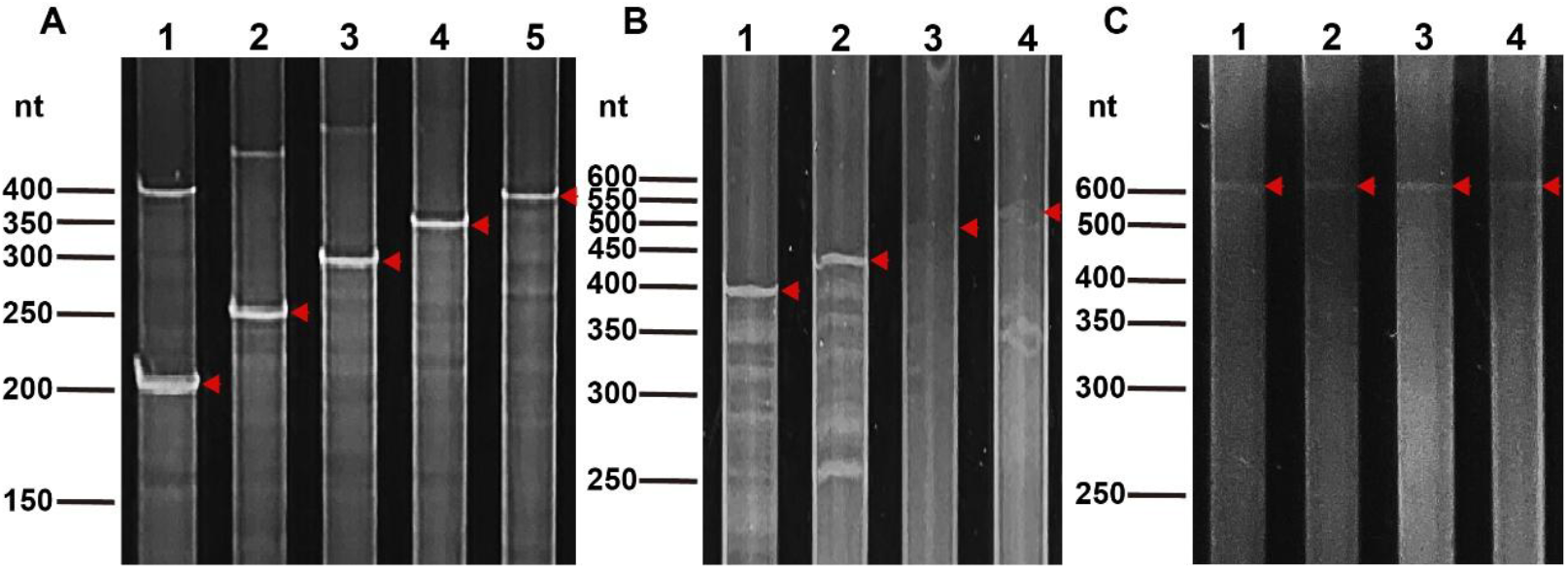
Validation of Synthesized ultra-long oligonucleotides. (A). We developed an innovative approach utilizing trimer phosphoramidites as building blocks for direct synthesis of extended DNA sequences on our engineered nonporous solid silica columns. (B). Polyacrylamide gel electrophoresis (PAGE) analysis of multiple poly(T) samples, each containing a single fragment of different length (200-400nt range). Lane 1: 200nt poly(T) oligo. Lane 2: 250nt poly(T) oligo. Lane 3: 300nt poly(T) oligo. Lane 4: 350nt poly(T) oligo. Lane 5: 400nt poly(T) oligo. (C). PAGE analysis of synthesized eSTR-containing fragments derived from the FRA10AC1 gene locus. Each lane represents a distinct fragment with varying lengths ranging from 400 to 650 nt. Lane 1: 400nt oligo. Lane 2: 451nt oligo with 17 repeats of eSTR locus. Lane 3: 499nt oligo with 33 repeats of eSTR locus. Lane 4: 550nt oligo with 50 repeats of eSTR locus. (D). PAGE analysis of synthesized 600nt fragments containing 270 CT repeats. Each lane represents the product synthesized by an individual synthesis column.

Short Tandem Repeats (STRs) are widespread genomic elements that significantly impact gene regulation^27^. Among these, expression STRs (eSTRs) represent a crucial subset that influences gene expression patterns. To further validate our synthesis platform, we randomly selected two eSTR loci with trinucleotide repeat units from previously identified eSTRs^28^. For each selected eSTR locus, we designed sequences incorporating 200 nucleotides upstream and downstream of the genomic region, with varying copy numbers of eSTR repeat units determining the final sequence length. Specifically, we designed four oligonucleotide sequences of different lengths, containing 0, 17, 33, and 50 repeat units (rpt), resulting in fragments of 400, 451, 499, and 550 nucleotides (nt), respectively. The designed sequences encompassed various challenging sequence types, including repetitive sequences, poly(T) sequences, and sequences with unbalanced base composition. Specifically, the APIP gene-associated eSTR locus featured AT-rich repetitive sequences and polyT sequences, while the FRA10AC1 gene-associated eSTR locus contained GC-rich repetitive sequences. These final 400-550nt oligonucleotides were synthesized using the method illustrated in Figure 1. PAGE analysis verified that all synthesized products exhibited their predicted molecular lengths (Figure 2B), demonstrating the high accuracy of our synthesis platform. We aimed to investigate whether the length of intrinsic repetitive sequences in synthetic fragments could exceed 400 nucleotides (nt). To this end, we designed a 600nt sequence comprising 270 consecutive cytosine-thymine (CT) repeats flanked by 30nt homologous sequences at each terminus. The sequence was independently synthesized using four separate synthesis columns. Polyacrylamide gel electrophoresis (PAGE) analysis confirmed successful synthesis of the target fragment with the expected molecular weight across all four replicates (Figure 2C). These results demonstrate the reproducibility and consistency of our synthetic platform in generating ultra-long challenging sequences.

### Selection and Validation of Synthetic DNA Sequences

Despite the high coupling efficiency of 99.5% achieved in chemical DNA synthesis^29^, the error accumulates progressively with increasing sequence length. For example, during the synthesis of a 300nt sequence, the proportion of error-free fragments in the final product is reduced to approximately 22%. Consequently, the major downstream challenge lies in isolating these error-free sequences from the complex mixture of synthetic products. To address this limitation and ensure sequence accuracy, we implemented a purification strategy utilizing biotinylated primers complementary to the 5’ termini of target sequences. Following hybridization, we isolated full-length products using streptavidin-functionalized magnetic beads and employed enzymatic mismatch repair to eliminate error-containing sequences (Figure 3A). The purified products were PCR-amplified and subcloned into appropriate vectors for subsequent analyses (Figure 3C). Agarose gel electrophoresis revealed distinct, expected sized bands for all synthetic sequences (Figure 3B), while Sanger sequencing confirmed sequence accuracy across all challenging motifs, regardless of length or complexity (Figure 3D).

**Figure 3.**
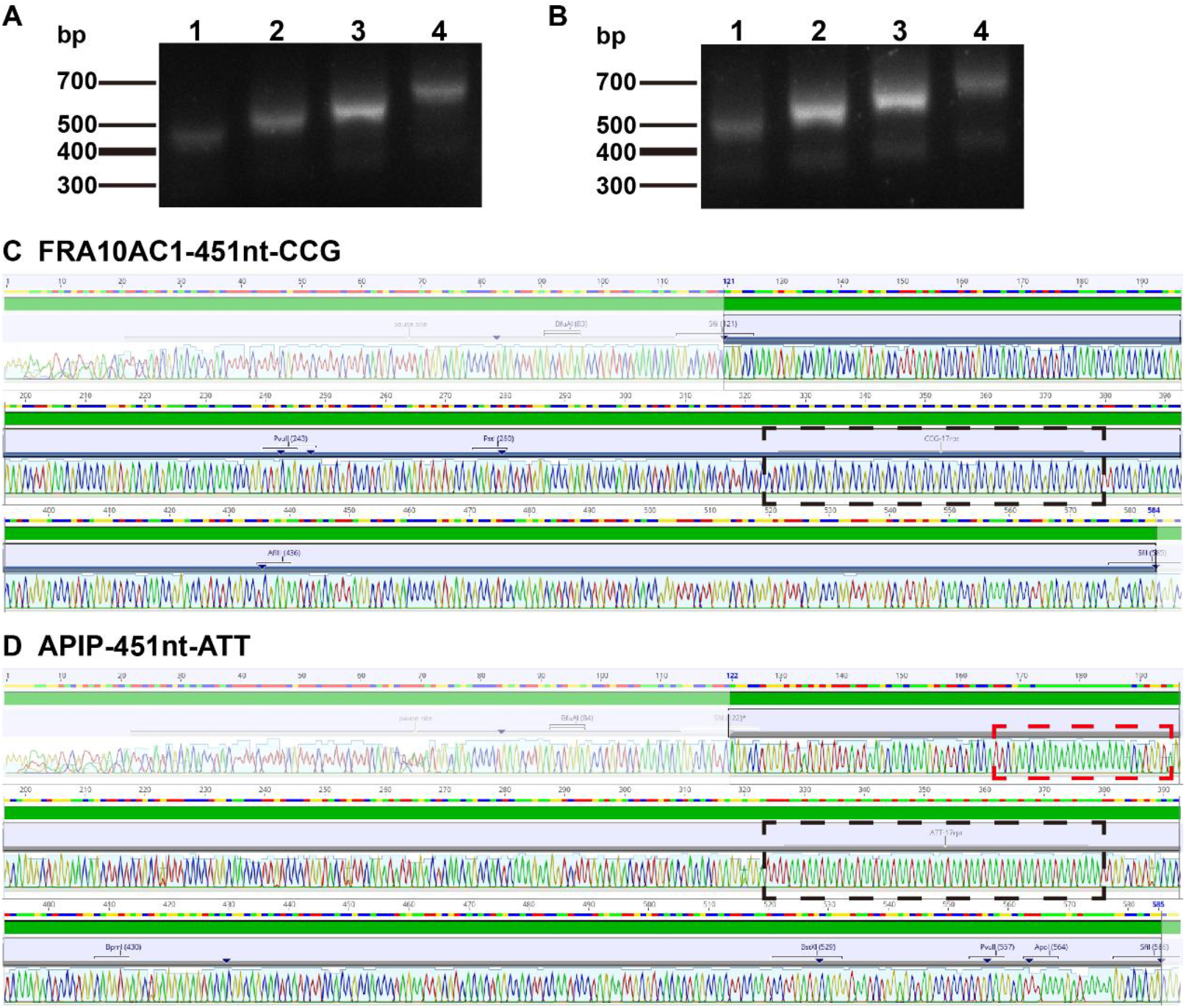
Quality control of selected and amplified oligonucleotides. (A). Agarose gel electrophoresis analysis of biotinylated primers-captured DNA fragments. Lanes 1-4 were 400 nt, 451 nt, 499 nt, and 550 nt in length, respectively. (B). Agarose gel electrophoresis analysis of mismatch repaired DNA fragments. Lanes 1-4 were 446 nt, 497 nt, 545 nt, and 596 nt in length, respectively, due to the primers containing homology arms required for subsequent Gibson assembly. (C-D). Sanger sequencing validation of synthetic FRA10AC1-related 451nt DNA sequences (C) and APIP-related 451nt DNA sequences (D). The black boxes indicate the repeat sequences (CCG or ATT) in each group, while the red box highlights the poly(T) sequences.

### Functional Validation of Synthetic Short Tandem Repeat Sequences

To validate the biological function of our synthetic sequences, we employed two reporter gene systems to investigate their impact on gene expression. Firstly, we focused on the EGFP reporter system (Figure 4A) to characterize the regulatory potential of our synthetic sequences. The backbone plasmid contained a minimal promoter (minP) driving the expression of enhanced green fluorescent protein (EGFP), which served as the baseline control for comparative analysis. We cloned 400nt and 451nt synthetic sequences upstream of the EGFP gene, positioned immediately before the minP into the backbone plasmid. These two sequence groups differed only in the presence or absence of the eSTR locus. Following transfection into HEK293T cells, fluorescence microscopy revealed that cells transfected with plasmids containing our synthetic sequences derived from the FRA10AC1 gene-associated eSTR loci exhibited significantly enhanced green fluorescent intensity compared to the control group. Notably, sequence with eSTR sites demonstrated a more pronounced upregulation of EGFP expression than the sequence without eSTR loci (Figure4 A). This enhanced EGFP expression suggests that our synthetic sequences possess robust regulatory capabilities in a cellular context. In contrast, the synthetic sequences derived from the APIP gene did not demonstrate a comparable upregulation effect (Figure 4A). This discrepancy may be attributed to the AT-rich features of the sequences. Recent studies reveal that the dinucleotide repeat motifs AT is depleted from enhancer sequences, suggesting that AT-rich sequences may play a non-essential or potentially inhibitory role in enhancer functionality^30^.

**Figure 4.**
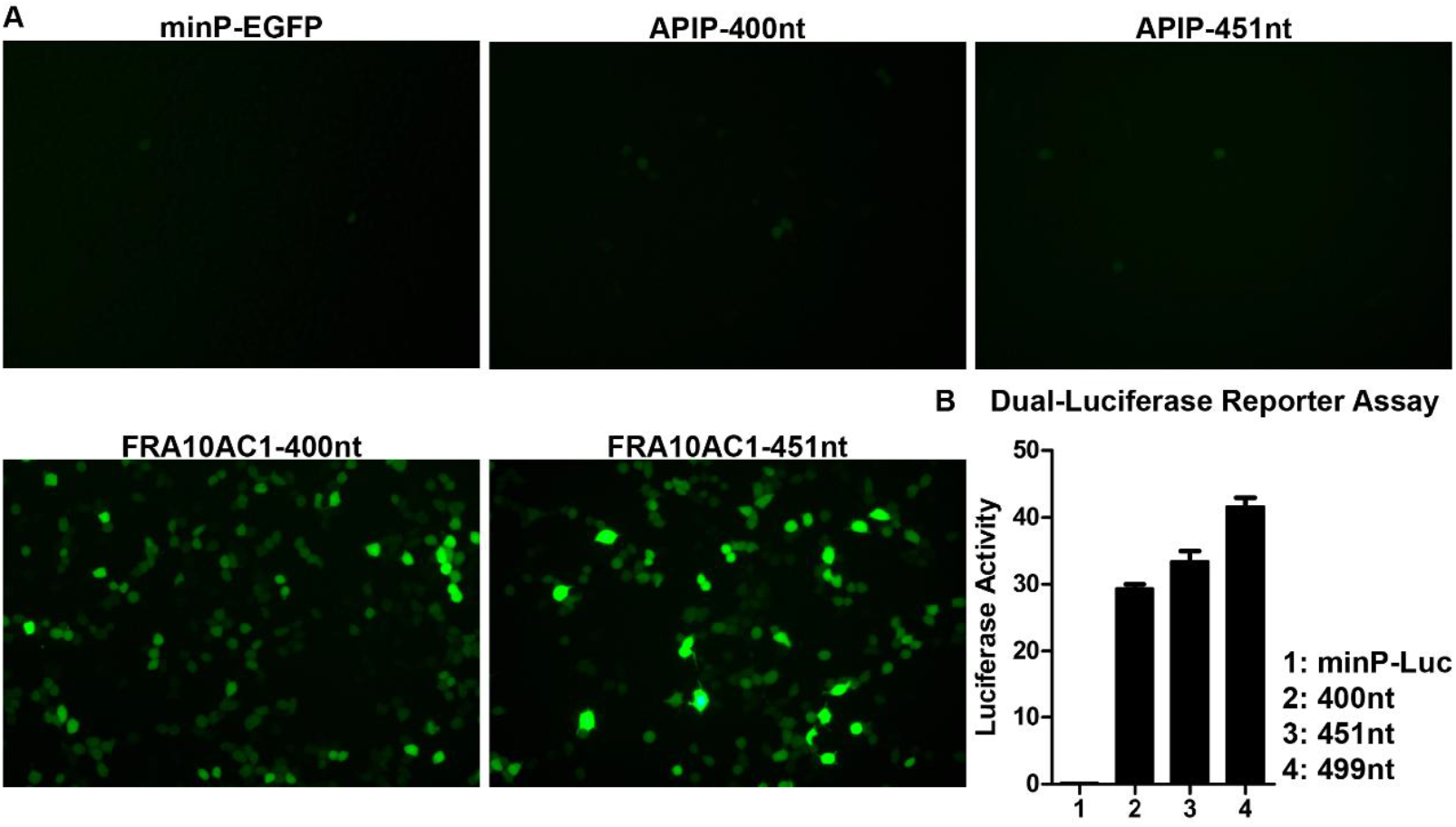
Functional validation of synthesized eSTR locus sequences. (A). Fluorescence microscopy imaging shows the effects of different eSTR loci on EGFP expression. APIP-associated loci showed no significant upregulation of downstream EGFP compared to the minimal promoter (minP). FRA10AC1-associated loci demonstrated substantial upregulation of EGFP expression. (B). Dual-luciferase reporter assays reveal a strong correlation between the repeat number of FRA10AC1-associated eSTR loci and downstream luciferase expression. Luciferase expression levels increased progressively with the increasing number of eSTR repeats.

Furthermore, we developed a dual-luciferase reporter system (Figure4 B) to systematically investigate the relationship between the FRA10AC1 gene-associated eSTR repeat numbers and downstream gene expression. We constructed plasmids incorporating 400nt, 451nt, and 499nt sequences upstream of the luciferase reporter gene, positioned before the minimal promoter (minP). These sequences corresponded to 0, 17, and 33 repeat units, respectively. The backbone plasmid without any synthetic sequences inserted served as the control, and Renilla luciferase was used as an internal control. Quantitative analysis of luciferase activities in transfected HEK293T cells demonstrated a significant positive correlation between eSTR repeat numbers and luciferase expression (Figure4 B). These results collectively indicate that our synthetic sequences not only maintain their regulatory function but also exhibit length-dependent modulation on downstream gene expression, with longer eSTR repeats conferring progressively stronger enhancement effects. The above functional analyses not only validated the accuracy of our synthetic sequences but also provided mechanistic insights into eSTR-mediated gene regulation.

### Precise Design and Validation of the Synthetic Biology Element

After successfully validating the effectiveness of our DNA synthesis platform in generating STR sequences within the 400-600 nt length range, we further explored its potential applications in synthesizing important elements of synthetic biology. Specifically, we selected a biologically significant target sequence for synthesis: the 5×HRE^31, 32^, which can be used to study cellular responses under hypoxic conditions. This element not only presents challenges in sequence complexity but also holds broad application prospects in synthetic biology.

The 5×HRE sequence consists of five repeated hypoxia-inducible factor binding sites (Figure 5A). In synthetic biology, the 5×HRE can be utilized to develop expression systems for specific genes under hypoxic conditions, thereby enabling the regulation of gene function in specific environments. We synthesized a 5×HRE sequence (Figure 5A) using the aforementioned DNA synthesis platform, which can be inserted upstream of specific gene sequences to induce expression in hypoxic environments. Sanger sequencing confirmed the precise synthesis (Figure 5B). The successful synthesis of the 5×HRE sequence not only demonstrates the technical advantages of our DNA synthesis platform but also provides new tools and methodologies for research in synthetic biology.

**Figure 5.**
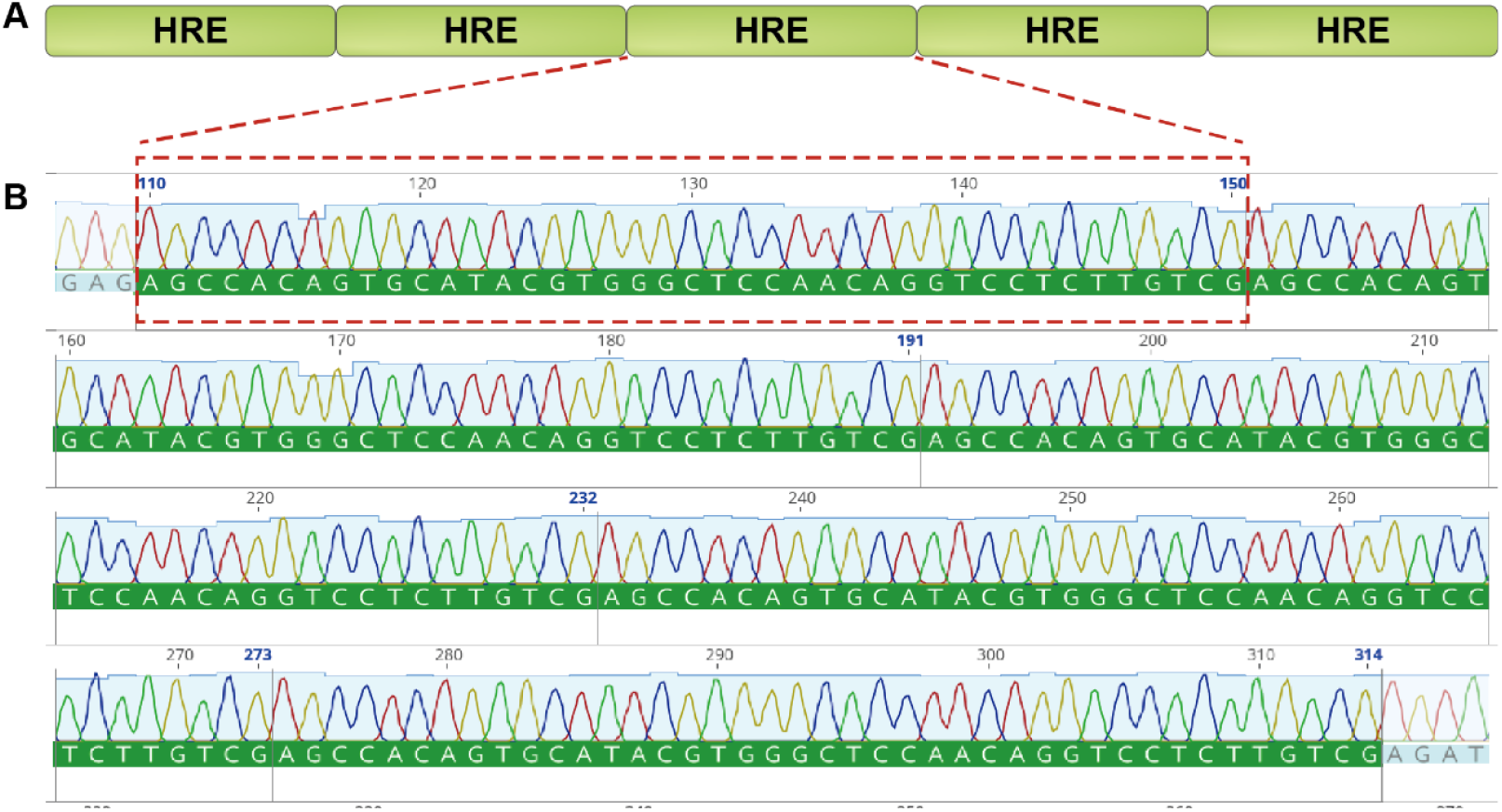
Validation of the synthesized 5×HRE sequence. (A). Schematic diagram of the 5×HRE sequence (B). Sanger sequencing validation of the synthetic 5×HRE sequence. Bases 110–150, 151–191, 192– 232, 233–273, and 274–314 each represent one of the five distinct HRE sequence frameworks.

## Discussion

The development of oligonucleotide synthesis technology holds significant importance for multiple fields, including synthetic biology and gene therapy. With the rapid advancement of life science technologies, DNA synthesis has become a critical technology driving continuous innovation. However, traditional DNA synthesis techniques are inherently limited in their capacity to synthesize long sequences exceeding 200 nucleotides and complex sequences, consequently constraining research progress in synthetic biology and precision medicine. Faced with increasingly complex biological research demands, breaking through existing synthesis technology limitations has become an urgent necessity. To address this challenge, we established an ultra-long sequence synthesis platform based on nonporous solid silica microspheres. We functionalized solid, non-porous silica microspheres and utilized these preprocessed microspheres to construct a synthesis column that served as the reaction vessel for DNA synthesis. This platform innovatively combines capture technology through biotin primers and enzymatic error correction strategies, successfully overcoming the length limitations of traditional oligonucleotide synthesis. Utilizing this platform, we successfully synthesized complex sequences up to 600 nucleotides in length. This breakthrough significantly expands the boundaries of oligonucleotide synthesis. Notably, the platform demonstrates exceptional capability in handling challenging complex sequences, including tandem repeat sequences, homopolymeric structures, and sequences with unbalanced base compositions.

The biotin primers-based capture technology and multi-step enzymatic error correction method represent key innovations of our platform. Long oligonucleotide synthesis primarily requires consideration of two aspects. First is synthesis efficiency. Compared to fragments under 200 nucleotides, the synthesis of longer oligonucleotides results in a significantly lower proportion of full-length fragments in the final product. A core technical challenge is purifying these low-quantity full-length fragments from the complex products. We designed a 20-nucleotide reverse complementary primer targeting the gene’s 5’ end, biotinylated at its 3’ terminus. When this biotin-labeled primer is annealed with synthesis products, it enables connecting successfully synthesized full-length fragments to streptavidin magnetic beads while removing non-full-length oligonucleotide fragments floating in the mixed system. Subsequently, low-cycle PCR amplification enriches these full-length fragments and releases the captured fragments from streptavidin magnetic beads. This step effectively improves the purity and accuracy of templates for subsequent enzymatic error correction, enhancing the efficiency of the error correction process and avoiding unexpected error fragments introduced by high template error rates.

Beyond full-length fragment enrichment and purification, long oligonucleotide synthesis must also consider synthesis accuracy. Previous literature reveals that common errors in oligonucleotide chemical synthesis include insertions, deletions, and substitutions, with corresponding error rates of 0.0045%, 0.1%, and 0.045%^33^. Consequently, during DNA chemical synthesis, the extension efficiency of each nucleotide incorporation is 99.8505% (100% - 0.0045% - 0.1% - 0.045%). When synthesizing a 400-nucleotide oligonucleotide, the theoretical final product accuracy is 54.97%. Actual accuracy may be even lower, with studies^34^ indicating that the actual accuracy of 400-nucleotide oligonucleotide synthesis is only 30%. Particularly when synthesizing G-base-rich sequences, the proportion of completely correct fragments significantly decreases. This is because G-base-induced errors occur more frequently than errors caused by other bases. For instance, G-base-induced substitution mutations occur at a frequency of 0.14% (dG to dA 0.11%, dG to dT 0.03%), representing the largest proportion of common substitution errors (total frequency 0.18%). We analyzed the accuracy of a 400-nucleotide GC-rich sequence (64% GC content, FRA10AC1-400nt) synthesized using our platform. Sanger sequencing results revealed a final product accuracy of 66.67%, far exceeding the theoretical accuracy. This result demonstrates that the multi-step enzymatic error correction step in our fragment preparation process is highly effective and crucial, significantly improving the proportion of completely correct final products and avoiding redundant cloning screening steps resulting from high error rates.

Direct synthesis of long oligonucleotide fragments can also conserve reagents consumed in full-length gene synthesis. Traditional gene synthesis involves fragmenting the full-length gene into approximately 80-nucleotide segments for synthesis and subsequent assembly (Figure 6A), requiring near-duplicate synthesis of nearly half the gene sequence to achieve effective assembly. In contrast, our synthesis platform requires each gene sequence nucleotide to be synthesized only once, avoiding redundant synthesis at junction points and dramatically reducing reagents and solvents used (Figure 6C). When synthesizing genes with repetitive sequences, conventional gene synthesis requires complex sequence fragmentation design and multiple synthesis steps (Figure 6B). Even for sequences under 500 nucleotides, 2-3 rounds of conventional gene synthesis are typically necessary. Our long-fragment synthesis platform, however, requires only a single synthesis cycle to obtain completely correct gene fragments or plasmids suitable for downstream applications (Figure 6C).

**Figure 6.**
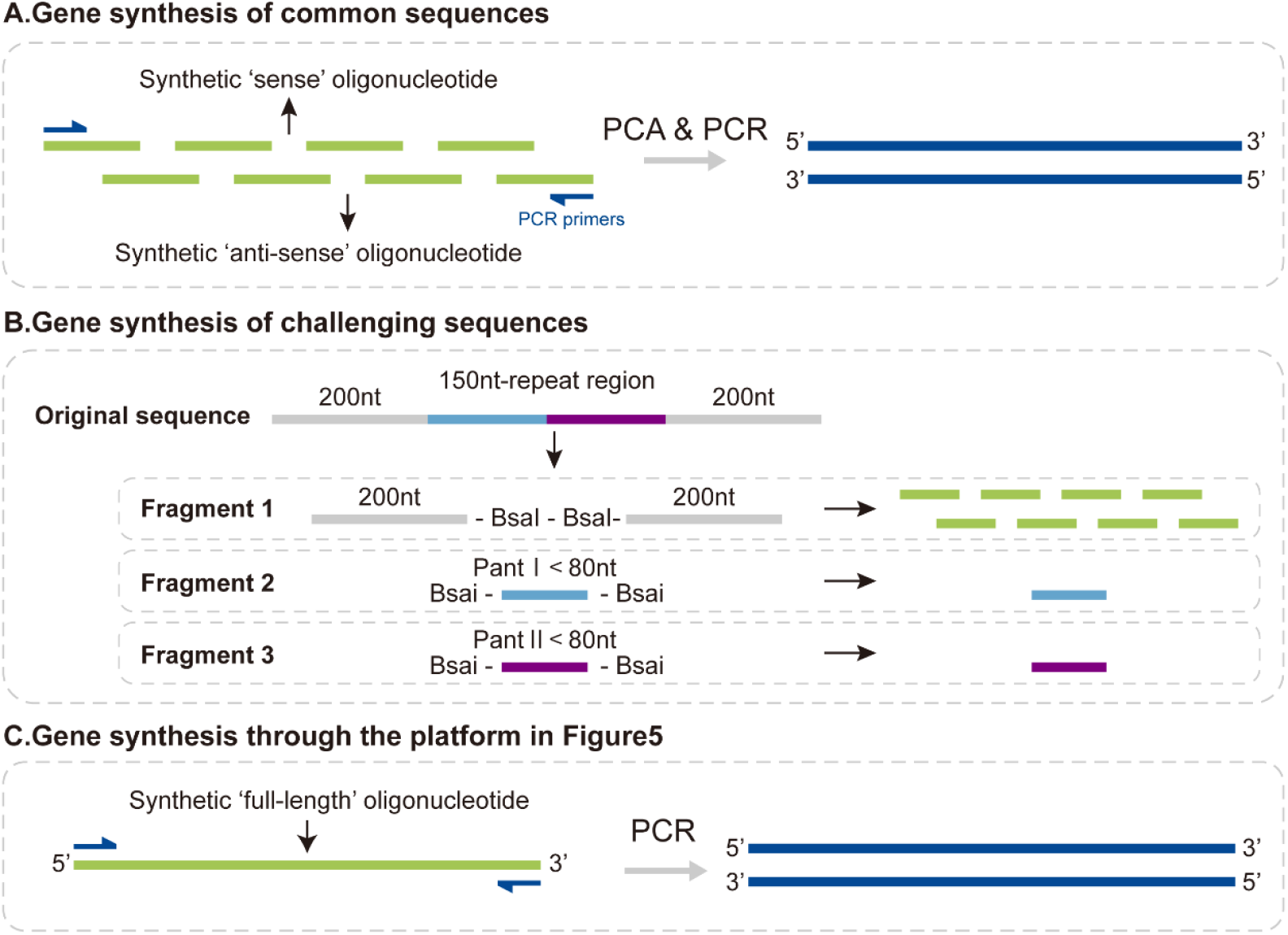
Comparison of conventional gene synthesis strategies and the long-fragment synthesis platform. (A). Schematic diagram of conventional gene synthesis for common sequences. The target sequence is divided into ~80 nt oligonucleotides with flanking sequences for bridging adjacent fragments. Synthesized sense and antisense sequences are assembled via polymerase cycling assembly (PCA). The complete synthetic fragment is then amplified by PCR. (B). Schematic diagram of gene synthesis for challenging sequences, exemplified by the 550 nt synthetic sequence in figure 2C. The full-length sequence is divided into two 200 nt flanking sequences (combined to form Fragment 1) and one 150 nt repetitive region (divided into Fragment 2 and Fragment 3). These three fragments are synthesized independently and correct fragments are ligated to obtain the full-length gene fragment. (C). Gene synthesis using the nonporous solid silica microsphere-based platform for long DNA fragments. Only a full-length sense sequence needs to be synthesized, which can then be PCR-amplified to obtain the complete synthetic fragment.

In addition to significantly reducing reagent consumption, our synthesis platform also effectively saves time costs. As illustrated in Figure 7, the detailed workflow and timeline of the UCOS synthesis platform are clear. For challenging sequences around 300 nucleotides in length, although our synthesis time in column is somewhat longer compared to conventional gene synthesis, the total time required to obtain a correct sequence suitable for downstream functional validation is as short as 50 hours. In contrast, traditional methods often require 2-3 rounds of gene synthesis to address the challenges associated with splicing, resulting in a overall timeline typically measured in weeks.

**Figure 7.**
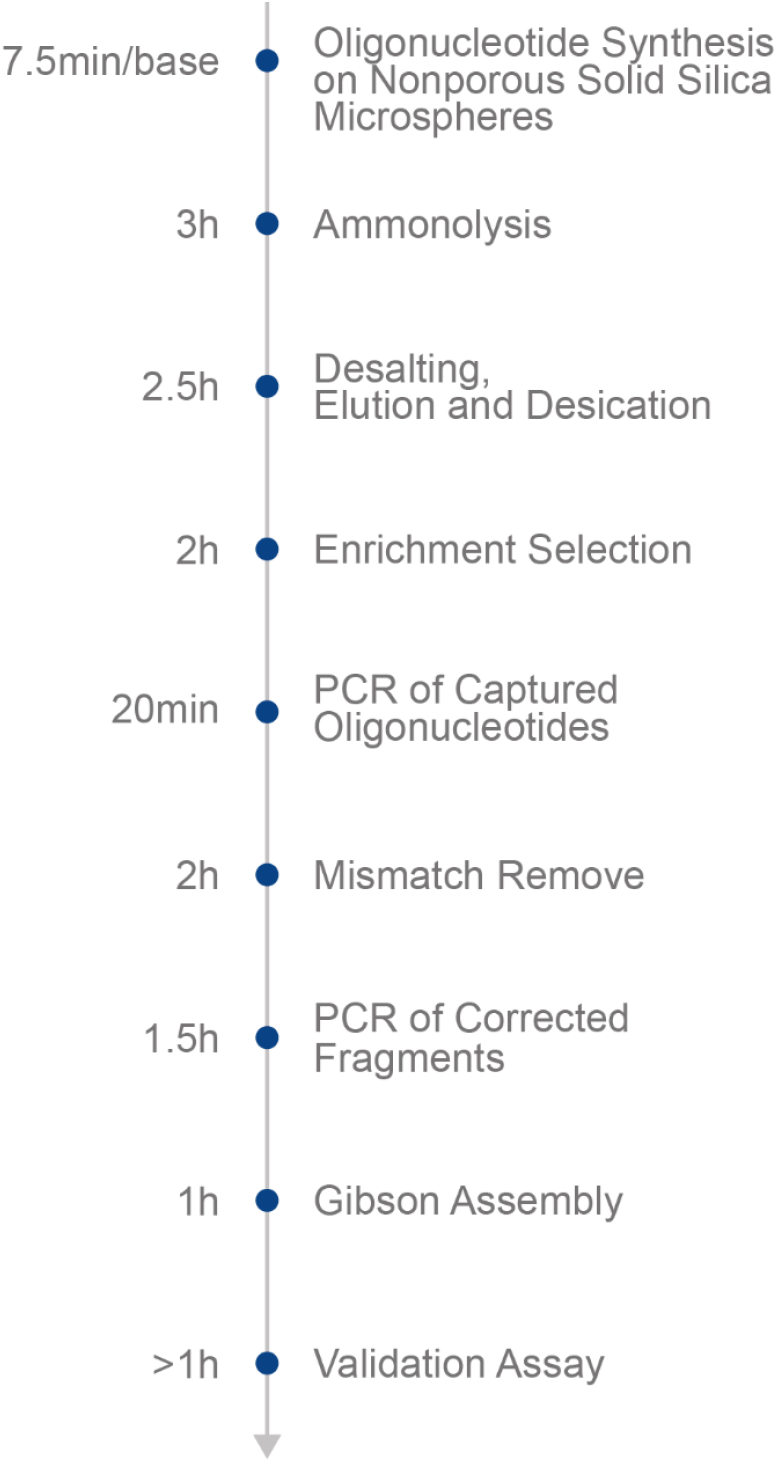
Detailed workflow and timeline for oligonucleotides synthesis on nonporous solid silica microspheres-based UCOS platform.

In summary, we have established a robust and practical workflow that encompasses long-fragment synthesis, accurate sequence selection, plasmid construction, and subsequent functional validation. In this study, we successfully employed this newly established synthetic platform to generate short tandem repeats (STRs) with repeat units of 1–6 nucleotides (Figure 8A). Nevertheless, this achievement does not signify the upper limit of the platform’s capabilities. A variety of naturally occurring yet highly complex sequences—many of which are difficult to synthesize via existing technologies—can also be synthesized using our platform. Notably, there are more tandem repeats in the genome with repeat units exceeding six nucleotides (Figure 8B), exhibiting high copy numbers. Our platform holds promise for advancing in-depth investigations of these loci at their native copy numbers. Moreover, the de novo design of some regulatory elements, such as 3’UTRs and circular RNAs, can require the synthesis of fully randomized sequences of 250 nt or longer (Figure 8C) for screening and validation^35^. Many protein-coding genes also harbor repetitive motifs that are important for synthetic biology. For instance, the pentapeptide sequence VPGVG—derived from elastin—contributes to elastin-like polypeptides^36^(Figure 8D), which alter their solubility with temperature changes and thus have broad potential applications in biomedicine and drug delivery. Similarly, synthesizing the repeat motifs of transcription activator–like effector (TALE) (Figure 8E) is especially challenging, as TALEs contain multiple repeats of a 34-amino-acid sequence^37^, with each repeat recognizing a specific DNA base through repeat variable di-residues (RVDs). Spider silk protein genes offer yet another example of naturally intricate protein-coding sequences. Their cDNA can span up to 2.4 kb and includes numerous repeats rich in glycine and alanine residues^38^ (Figure 8F), with the longest repeat unit reaching 34 amino acids (equivalent to 102 nucleotides). Taken together, further refinements in our platform for synthesizing ultra-long and complex DNA sequences may enable rapid and reliable assembly of these challenging constructs, thereby opening new avenues for both fundamental biomedical research and a wide range of practical applications.

**Figure 8.**
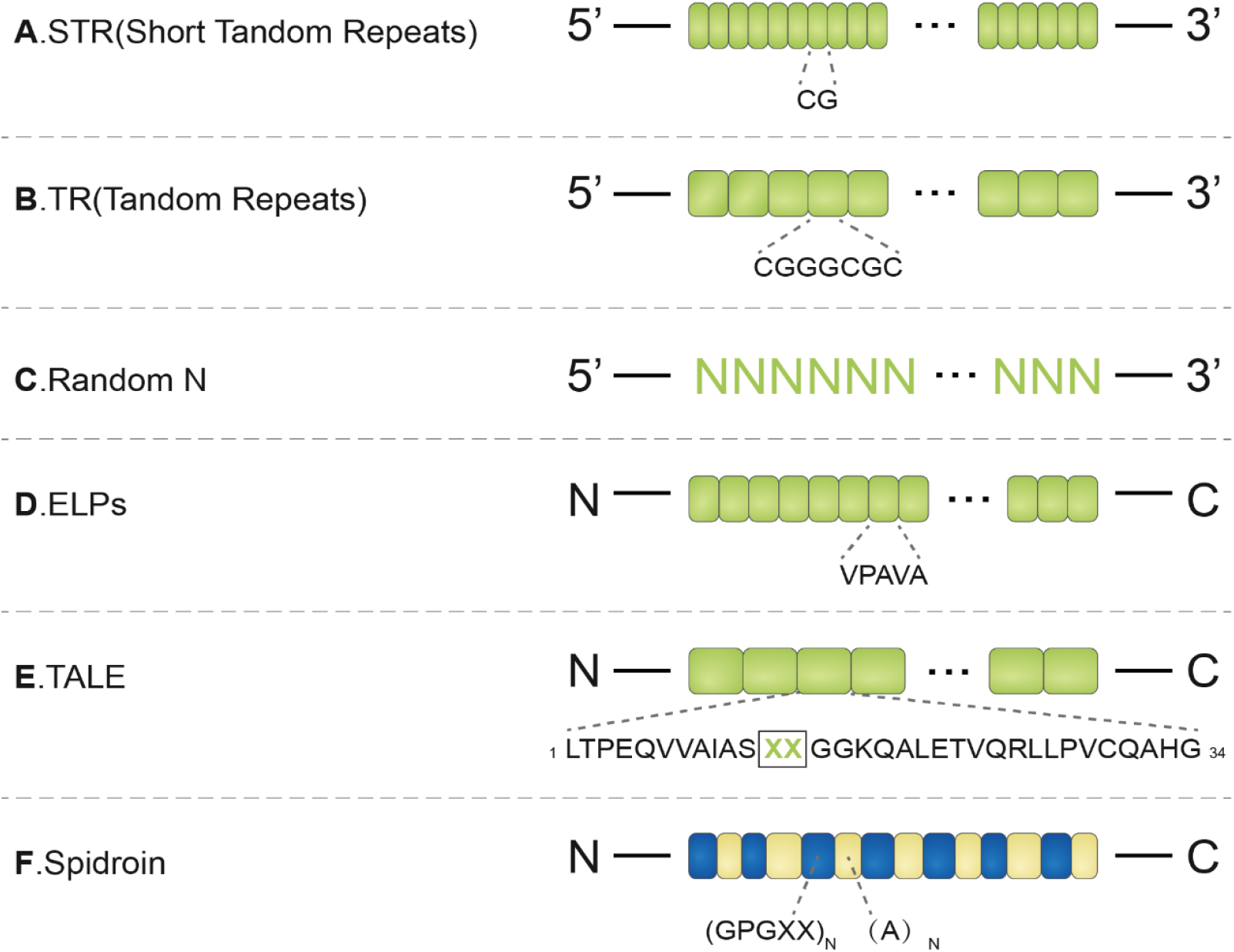
Schematic illustration of potential complex and challenging sequences that can be synthesized using our platform. (A). Short tandem repeat sequences. The repeat unit is in the 1–6 nt range and typically exhibits an imbalanced base composition. (B). Tandem repeat sequences. The repeat unit is longer than 6 nt and generally also exhibits an imbalanced base composition. (C). Long random sequences. De novo design of regulatory elements such as circular RNAs or 3’UTRs often requires the synthesis of fully random sequences of 250 nt or more. (D). Elastin-like polypeptides. The temperature sensitivity of these proteins depends primarily on longer sequences formed by tandem repeats of a pentapeptide motif. (E). Transcription activator-like effectors (TALEs). The core region of these proteins comprises tandem repeats of a 34-amino-acid sequence that is nearly identical. The specific nucleotide (A, T, C, or G) recognized by TALE is determined by the 12th and 13th residues, known as repeat variable di-residues (RVDs). (F). Spider silk proteins. These natural protein-coding genes are enriched in glycine and alanine residues and contain numerous tandem repeats.

## Methods

### Ultralong oligonucleotides synthesis

Our novel synthesis platform is based on nonporous solid silica microspheres for long-sequence DNA synthesis. The optimized preparation protocol for nonporous solid silica microspheres begins with pretreatment of nonporous solid silica microspheres using 2M ammonium fluoride solution to generate hydroxyl-rich surfaces. Subsequently, a primary grafting step was performed using 10% (w/w) GPOS silane reagent in acetonitrile. The modified microspheres then underwent a secondary grafting process with DMT-diamine in the presence of alkylamine DMF, followed by deprotection using trichloroacetic acid to remove protecting groups. The final functionalization step involved a condensation reaction between the modified microspheres and linker molecules, Unylinker, in the presence of condensing agents (DMSO, DIPEA, and HBTU). Notably, to prevent unwanted reactions between hydroxyl groups located at the intersection of primary and secondary grafting reagents and downstream linker molecules, we performed capping of these groups using acetic anhydride. This comprehensive surface modification protocol yielded nonporous solid silica microsphere supports suitable for oligonucleotide synthesis. Significantly, we discovered that utilizing amine groups as active functional moieties, rather than hydroxyl groups, substantially improved the achievable length of ultra-long oligonucleotides. Then we employ the standard phosphoramidite methodology for the synthesis of ultralong oligonucleotides on the DYHOW2B synthesis equipment(Dynegene Technologies) with our nonporous solid silica microspheres supports columns.

### Validation of Ultralong oligonucleotides

We designed synthetic sequences based on genomic regions encompassing 200 base pairs upstream and downstream of selected expression Short Tandem Repeat (eSTR) loci. The final synthetic sequences were constructed by inserting eSTR repeats (0, 17, 33, and 50 repeat units) into the central region of the basic sequence. Synthesis was performed using phosphoramidite trimer chemistry with Silica microsphere-packed synthesis columns prepared according to the method illustrated in Ultralong oligonucleotides synthesis. Synthesized sequences underwent desalting purification. The purity and fragment size of the synthesis products were verified using Polyacrylamide Gel Electrophoresis (PAGE). A 8% polyacrylamide gel containing 7 M urea was prepared in 1× TBE buffer. The gel solution was filtered through a 0.22 μm membrane and degassed for 15 minutes. Polymerization was initiated by adding 0.1% (w/v) ammonium persulfate (APS) and 0.01% (v/v) N,N,N’,N’-tetramethylethylenediamine (TEMED). DNA samples (2-4 μL, 100 ng/μL) were mixed with an equal volume of 2× loading buffer (#AM8546G,Invitrogen, USA) and denatured at 100°C for 5 minutes, followed by immediate cooling on ice. A 50bp DNA ladder (#ALH313, Balb, China) was used as a size marker. Electrophoresis was performed at 200V constant power for 40 minutes in 1× TBE buffer at room temperature. The gel was stained with SYBR Gold (1:10,000 dilution in 1× TBE; #S11494,Invitrogen, USA) for 10-15 minutes in the dark with gentle agitation. After three washes with deionized water, the gel was visualized using a BluSight Pro system (#GD50502, Monad, China). Detailed information for the two eSTR loci synthetic sequences is provided in Supplementary Table 1.

### Enrichment of full-length oligonucleotides

A complementary primer was designed targeting the 5’ end of each synthetic sequence, with biotin conjugated to the primer’s 3’ terminus. Primers and synthetic fragments were diluted in IDT Duplex buffer at equimolar ratios and subjected to the following annealing protocol: heating to 98° C, then gradually cooling to 60°C at a rate of 1°C per minute. Incubating at 64°C for 10 min, then gradually cooling to room temperature at a rate of 1°C per minute. Biotinylated primers and their corresponding synthetic fragments were purified using QuarAcces Hyper Enrichment Beads (#ND3018, Dynegene, China).

### Mismatch remove of full-length oligonucleotides

The purified fragments were amplified by PCR using 2× PCR Mix (#NL4001, Dynegene, China) with specific primers for each eSTR. PCR amplification was performed under the following conditions: initial denaturation at 95°C for 30 s, followed by 10 cycles of denaturation at 95°C for 15 s, annealing at 60°C for 15 s, and extension at 72°C for 1 min. A final extension step was conducted at 72°C for 3 min. Then, the purified amplification products were prepared for error correction by adding 1 µL of 10X Reaction Buffer I (#NE1003, Dynegene, China) and nuclease-free water to a total volume of 10 µL, with initial PCR thermal cycling performed at 98°C for 2 min, followed by sequential steps at 4°C for 5 min, 37°C for 5 min, and a final hold at 4°C. After reaching the final hold step, 2 µL of RecJf Exonuclease (#NE1003, Dynegene, China) and 6 µL of CorrectASE I (#NE1003, Dynegene, China) were added to the annealed product. The mixture was then incubated at 37°C for 1 hour, with the thermal cycler lid temperature set to 47°C. Following the previous reaction, a 1 µL aliquot of 100-fold diluted CorrectASE II was added. The sample was then incubated at 37°C for 30 minutes, with the thermal cycler lid temperature maintained at 47°C.

### Complex Gene Construction

The error-corrected product was then prepared for PCR amplification by adding fragment-specific primers (10 µM, 2 µL each), 25 µL of 2× PCR Mix (#NL4001, Dynegene, China), and nuclease-free water to a total volume of 50 µL. PCR amplification was conducted using the following thermal cycling conditions: an initial denaturation at 95°C for 3 min, followed by 25 cycles of denaturation at 95°C for 15 sec, annealing at 60°C for 15 sec, and extension at 72°C for 1 min, with a final extension step at 72°C for 5 min and a terminal hold at 4°C. The amplified error-corrected product was purified with QuarAcces Hyper Pure Beads (#ND3011, Dynegene, China) and eluted with nuclease-free water and quantified using a Thermo NanoDrop OneC. It is worth noting that for sequences with high GC content, 5% DMSO was supplemented to the PCR reaction mixture. Detailed primer sequence information is provided in Supplementary Table 2. All primers were synthesized by Dynegene Technologies.

For 5×HRE sequence, Gibson Assembly(#NG1002, Dynegene, China) was employed to insert the error-corrected fragments into the linearized vector carrying a complete downstream expression cassette. Sanger sequencing was used to assess the complete accuracy of the final sequence.

### Functional verification of Complex Gene Construction based on Ultralong oligonucleotides synthesis

We constructed a series of plasmids based on the pGL4 vector backbone, which were synthesized by Dynegene Technologies according to the sequence information available on the Addgene homepage. The plasmid minP-EGFP was generated by inserting a minimal promoter (minP) sequence and the EGFP reporter gene, with two SfiI restriction sites featuring different sticky ends inserted upstream of the minP. The linearized minP-EGFP plasmid was prepared using SfiI restriction enzyme (#R0123S, New England Biolabs, USA). Gibson Assembly(#NG1002, Dynegene, China) was employed to insert the error-corrected eSTR fragments into the linearized vector, using primers with homology arms corresponding to the vector’s SfiI flanking sequences. Plasmids were extracted using a endotoxin-free plasmid extraction kit (#12163, QIAGEN, Germany) and transfected into HEK293T cells using polyethyleneimine (PEI; #24765-1, PolySciences, USA). EGFP expression was visualized via fluorescence microscopy 24 hours post-transfection.

We replaced the EGFP reporter gene with Luciferase in the previously constructed plasmids. A Renilla luciferase-expressing plasmid (#E6911, Promega, USA) was used as an internal control. HEK293T cells were co-transfected with Luciferase and Renilla plasmids at a 50:1 ratio. Dual-luciferase activity was measured using a luminometer 24 hours post-transfection, following the detailed protocol provided in the Promega dual-luciferase reporter assay kit (#E1910, Promega, USA).

## Acknowledgment

This work was supported by Shanghai Municipal Science and Technology Major Project, the National Key R&D Program of China (2023YFA0913804, 2024YFA0916603, 2022FYC2503300), Shanghai Science and Technology Innovation Action Program (No. 24JS2840400, 24ZR1439900), Shanghai Municipal Health Commission Collaborative Innovation Group(2024CXJQ03), Natural Science Foundation of China (32370724, 82401759), Shanghai Municipal Commission of Education(2024AIZD016), the fundamental research funds for the central universities (YG2021ZD2020, YG2023QNB12, YG2023QNB20, YG2021QN143).

## Author contributions

Mancang Zhang, Yongyong Shi: Conceptualization, Methodology. Mancang Zhang, Yang Hu: Investigation, Validation, Writing-Original draft. Yang Hu, Hao Huang: Software, Data curation, Formal analysis. Yongyong Shi: Supervision, Writing-Reviewing and Editing.

## Data Availability Statement

The data that support the findings of this study are available from the corresponding author upon reasonable request. The scripts used in this manuscript are listed in the supplemental material.

## References

1. Elworth, R.; Diaz, C.; Yang, J.; Figueiredo, P.; Ternus, K.; Treangen, T., Synthetic DNA and biosecurity: Nuances of predicting pathogenicity and the impetus for novel computational approaches for screening oligonucleotides. PLOS Pathogens 2020, 16, e1008649.

2. Weber, W.; Fussenegger, M., The impact of synthetic biology on drug discovery. Drug discovery today 2009, 14, 956–63.

3. Yew, N.; Zhao, H.; Przybylska, M.; Wu, I. H.; Tousignant, J.; Scheule, R.; Cheng, S. H., CpG-Depleted Plasmid DNA Vectors with Enhanced Safety and Long-Term Gene Expression in Vivo. Molecular therapy : the journal of the American Society of Gene Therapy 2002, 5, 731–8.

4. Michelson, A.; Todd, A., Nucleotides part XXXII. Synthesis of a dithymidine dinucleotide containing a 3′: 5′-internucleotidic linkage. Journal of The Chemical Society (resumed) 1955, 1955.

5. Khorana, H.; Razzell, W.; Gilham, P.; Tener, G.; Pol, E., SYNTHESES of dideoxyribonucleotides. Journal of the American Chemical Society 1957, 79.

6. Beaucage, S.; Caruthers, M. H., Deoxynucleoside phosphoramidites—A new class of key intermediates for deoxypolynucleotide synthesis. Tetrahedron Letters 1981, 22, 1859–1862.

7. Matteucci, M.; Caruthers, M., ChemInform Abstract: SYNTHESIS OF DEOXYOLIGONUCLEOTIDES ON A POLYMER SUPPORT. Chemischer Informationsdienst 1981, 12.

8. Hoose, A.; Vellacott, R.; Storch, M.; Freemont, P.; Ryadnov, M., DNA synthesis technologies to close the gene writing gap. Nature Reviews Chemistry 2023, 7, 144–161.

9. Mani, I., Recent development in DNA synthesis technology. 2022; pp 1–8.

10. Hölz, K.; Hoi, J.; Schaudy, E.; Somoza, V.; Lietard, J.; Somoza, M., High-Efficiency Reverse (5′→3′) Synthesis of Complex DNA Microarrays. Scientific Reports 2018, 8.

11. Kosuri, S.; Church, G., Large-scale de novo DNA synthesis: Technologies and applications. Nature methods 2014, 11, 499–507.

12. Stanley, P.; Vickers, A.; Strittmatter, L.; Lee, K., Decoding DNA data storage for investment. Biotechnology Advances 2020, 45, 107639.

13. Stemmer, W.; Crameri, A.; Ha, K.; Brennan, T.; Heyneker, H., Stemmer, W. P. C., Crameri, A., Ha, K. D., Brennan, T. M. & Heyneker, H. L. Single-step PCR assembly of a gene and a whole plasmid from large numbers of oligonucleotides. Gene 164, 49-53. Gene 1995, 164, 49–53.

14. TerMaat, J.; Pienaar, E.; Whitney, S.; Mammedov, T.; Subramanian, A., Gene synthesis by integrated polymerase chain assembly and PCR amplification using a high-speed thermocycler. Journal of microbiological methods 2009, 79, 295–300.

15. Gibson, D., Enzymatic assembly of DNA molecules up to several hundred kilobases. Protocol Exchange 2009, 6.

16. Gibson, D.; Benders, G.; Andrews-Pfannkoch, C.; Denisova, E.; Baden-Tillson, H.; Zaveri, J.; Stockwell, T.; Brownley, A.; Thomas, D.; Algire, M.; Merryman, C.; Young, L.; Noskov, V.; Glass, J.; Venter, J.; Hutchison, C.; Smith, H., Complete Chemical Synthesis, Assembly, and Cloning of a Mycoplasma genitalium Genome. Science (New York, N.Y.) 2008, 319, 1215–20.

17. Gibson, D.; Smith, H.; Hutchison, C.; Venter, J.; Merryman, C., Chemical synthesis of the mouse mitochondrial genome. Nature methods 2010, 7, 901–3.

18. Bollum, F., Thermal Conversion of Nonpriming Deoxyribonucleic Acid to Primer. The Journal of biological chemistry 1959, 234, 2733–4.

19. Schott, H.; Schrade, H., Single-step elongation of oligodeoxynucleotides using terminal deoxynucleotidyl transferase. European journal of biochemistry / FEBS 1984, 143, 613–20.

20. Lu, X.; Li, J.; Li, C.; Lou, Q.; Peng, K.; Cai, B.; Liu, Y.; Yao, Y.; Lu, L.; Tian, Z.; Ma, H.; Wang, W.; Cheng, J.; Guo, X.; Jiang, H.; Ma, Y., Enzymatic DNA Synthesis by Engineering Terminal Deoxynucleotidyl Transferase. ACS Catalysis 2022, 12 (5), 2988–2997.

21. Hamilton, W.; Claessens, A.; Otto, T.; Fairhurst, R.; Rayner, J.; Kwiatkowski, D., Extreme mutation bias and high AT content in Plasmodium falciparum. Nucleic acids research 2016, 45.

22. Scheller, J.; Guehrs, K.-H.; Grosse, F.; Conrad, U., Production of spider silk proteins in tobacco and potato. Nature biotechnology 2001, 19, 573-7.

23. Willems, T.; Gymrek, M.; Highnam, G.; Mittelman, D.; Erlich, Y., The landscape of human STR variation. Genome research 2014, 24.

24. Hirashima, K.; Seimiya, H., Telomeric repeat-containing RNA/G-quadruplex-forming sequences cause genome-wide alteration of gene expression in human cancer cells in vivo. Nucleic acids research 2015, 43.

25. Ennifar, E.; Nikulin, A.; Tishchenko, S.; Serganov, A.; Nevskaya, N.; Garber, M.; Ehresmann, B.; Ehresmann, C.; Nikonov, S.; Dumas, P., The crystal structure of UUCG tetraloop. Journal of molecular biology 2000, 304, 35-42.

26. Aarons, S.; Abbas, A.; Adams, C.; Fenton, A.; O’Gara, F., A Regulatory RNA (PrrB RNA) Modulates Expression of Secondary Metabolite Genes in Pseudomonas fluorescens F113. Journal of bacteriology 2000, 182, 3913-9.

27. Gemayel, R.; Vinces, M.; Legendre, M.; Verstrepen, K., Variable Tandem Repeats Accelerate Evolution of Coding and Regulatory Sequences. Annual review of genetics 2010, 44, 445-77.

28. Fotsing, S. F.; Margoliash, J.; Wang, C.; Saini, S.; Yanicky, R.; Shleizer-Burko, S.; Goren, A.; Gymrek, M., The impact of short tandem repeat variation on gene expression. Nature Genetics 2019, 51 (11), 1652-1659.

29. Hughes, R.; Ellington, A., Synthetic DNA Synthesis and Assembly: Putting the Synthetic in Synthetic Biology. Cold Spring Harbor Perspectives in Biology 2017, 9, a023812.

30. Yáñez-Cuna, J. O.; Arnold, C. D.; Stampfel, G.; Boryń, Ł. M.; Gerlach, D.; Rath, M.; Stark, A., Dissection of thousands of cell type-specific enhancers identifies dinucleotide repeat motifs as general enhancer features. Genome Research 2014, 24 (7), 1147-1156.

31. Semenza, G.; Nejfelt, M.; Chi, S.; Antonarakis, S., Hypoxia-inducible nuclear factors bind to an enhancer element located 3Â to the human erythropoietin gene. Proceedings of the National Academy of Sciences of the United States of America 1991, 88, 5680-4.

32. Harris, A. L., Hypoxia — a key regulatory factor in tumour growth. Nature Reviews Cancer 2002, 2 (1), 38-47.

33. Masaki, Y.; Onishi, Y.; Seio, K., Quantification of synthetic errors during chemical synthesis of DNA and its suppression by non-canonical nucleosides. Scientific Reports 2022, 12 (1).

34. Yin, Y.; Arneson, R.; Apostle, A.; Eriyagama, A. M. D. N.; Chillar, K.; Burke, E.; Jahfetson, M.; Yuan, Y.; Fang, S., Long oligodeoxynucleotides: chemical synthesis, isolation via catching-by-polymerization, verification via sequencing, and gene expression demonstration. Beilstein Journal of Organic Chemistry 2023, 19, 1957-1965.

35. Liu, L.-H.; Chen, J.; Lai, S.; Zhao, X.; Yang, M.; Wu, Y.-R.; Zhang, Z.; Jiang, A., Functional RNA mining using random high-throughput screening. Nucleic Acids Research 2025, 53 (2).

36. Fletcher, E.; Yan, D.; Kosiba, A.; Zhou, Y.; Shi, H., Biotechnological applications of elastin-like polypeptides and the inverse transition cycle in the pharmaceutical industry. Protein Expression and Purification 2018, 153.

37. Boch, J.; Scholze, H.; Schornack, S.; Landgraf, A.; Hahn, S.; Thieme, S.; Lahaye, T.; Nickstadt, A.; Bonas, U., Breaking the code of DNA binding specificity of TAL-Type III effectors. Science (New York, N.Y.) 2009, 326, 1509-12.

38. Spiess, K.; Lammel, A.; Scheibel, T., Recombinant Spider Silk Proteins for Applications in Biomaterials. Macromolecular Bioscience 2010, 10 (9), 998–1007.

