## Supplementary Table 1 for "Chemically Synthesized Ultra-long DNA as Building Blocks to Accelerate Complex Gene Construction in Synthetic Biology"

| Gene ID | Sequences |
|---|---|
| APIP-400nt | AGGGAGATAGAGAAATAATTGCTTCTTTGATAACATCCTAACGTTCTTTTTTTTTTTTTTTTCTAGGATGGCTGGGCAGGGTTCTGCAACACAGCAGAAACCCCAGGCTTCCCAAGACCCAGGATCAGAACTGTATAATGTCACTTCTGCCTCACTTTGTTGGACAAAGTATGTCACAAGATCATCTCAGTGGTGGTAAATTAAGGCAGAGTCTCACTTTGTCACCCAGGCTGGAGTGCAGTGGTGTGATCTCGGCTGCTCACTACAACCTCCGCCTCCCATGTTAAGCGACTCTCATGCCTCAACCTTACCCATGTTAAGCGACTCTCATGCCTCAACCTTACCAAGTAGCTGGGATTACAGGTGCATGCCACCACAGCTGGCTAAATTTTGTATTTTT |
| APIP-451nt | AGGGAGATAGAGAAATAATTGCTTCTTTGATAACATCCTAACGTTCTTTTTTTTTTTTTTTTCTAGGATGGCTGGGCAGGGTTCTGCAACACAGCAGAAACCCCAGGCTTCCCAAGACCCAGGATCAGAACTGTATAATGTCACTTCTGCCTCACTTTGTTGGACAAAGTATGTCACAAGATCATCTCAGTGGTGGTAAAAttattattattattattattattattattattattattattattattattattTTAAGGCAGAGTCTCACTTTGTCACCCAGGCTGGAGTGCAGTGGTGTGATCTCGGCTGCTCACTACAACCTCCGCCTCCCATGTTAAGCGACTCTCATGCCTCAACCTTACCCATGTTAAGCGACTCTCATGCCTCAACCTTACCAAGTAGCTGGGATTACAGGTGCATGCCACCACAGCTGGCTAAATTTTGTATTTTT |
| APIP-499nt | AGGGAGATAGAGAAATAATTGCTTCTTTGATAACATCCTAACGTTCTTTTTTTTTTTTTTTTCTAGGATGGCTGGGCAGGGTTCTGCAACACAGCAGAAACCCCAGGCTTCCCAAGACCCAGGATCAGAACTGTATAATGTCACTTCTGCCTCACTTTGTTGGACAAAGTATGTCACAAGATCATCTCAGTGGTGGTAAAAttattattattattattattattattattattattattattattattattattattattattattattattattattattattattattattattattattattTTAAGGCAGAGTCTCACTTTGTCACCCAGGCTGGAGTGCAGTGGTGTGATCTCGGCTGCTCACTACAACCTCCGCCTCCCATGTTAAGCGACTCTCATGCCTCAACCTTACCCATGTTAAGCGACTCTCATGCCTCAACCTTACCAAGTAGCTGGGATTACAGGTGCATGCCACCACAGCTGGCTAAATTTTGTATTTTT |
| APIP-550nt | AGGGAGATAGAGAAATAATTGCTTCTTTGATAACATCCTAACGTTCTTTTTTTTTTTTTTTTCTAGGATGGCTGGGCAGGGTTCTGCAACACAGCAGAAACCCCAGGCTTCCCAAGACCCAGGATCAGAACTGTATAATGTCACTTCTGCCTCACTTTGTTGGACAAAGTATGTCACAAGATCATCTCAGTGGTGGTAAAAttattattattattattattattattattattattattattattattattattattattattattattattattattattattattattattattattattattattattattattattattattattattattattattattattattattattattattattattattattattTTAAGGCAGAGTCTCACTTTGTCACCCAGGCTGGAGTGCAGTGGTGTGATCTCGGCTGCTCACTACAACCTCCGCCTCCCATGTTAAGCGACTCTCATGCCTCAACCTTACCCATGTTAAGCGACTCTCATGCCTCAACCTTACCAAGTAGCTGGGATTACAGGTGCATGCCACCACAGCTGGCTAAATTTTGTATTTTT |
| FRA10-400nt | TTTGAGGTCCTTCTCCCTGATGCCAAATATACCCACAGCCCTCCTCACACACCCTTTCCCTTCCTCCCTGTGTGCCAAGTTCGCCCCCGGAGTCGTCGTTTCCTTCTTTCCCGGCAGCTGCAGCGACGACCCACGGCCTGAGAGAGCCGCTGCAGCACAGGTCCCGTGCGCCCTGCCGCACAGCCTCGCCACAACCACCACGCAACCCGCCTCTCCCTACGGGTCCCGACTGGGCACCACTTC |

| | |
|---|---|
| | CGGTCCGACACGGCCACGTGTTACATCTAAATGGCACCGTCCCCCGAGT<br>GCGCCGACCTTGTGCTACAGCCAGAACAGCTCCGGGAAACAAGGGCA<br>GCGGGAGAAACGGCCCGGAAAGGGACGAAGAGGCTTCCAGCTTTCGC<br>CAAGTCGCGGCTCC |
| FRA10-451nt | TTTGAGGTCCTTCTCCCTGATGCCAAATATACCCACAGCCCTCCTCACAC<br>ACCCTTTCCCTTCCTCCCTGTGTGCCAAGTTCGCCCCCGGAGTCGTCGT<br>TTCCTTCTTTCCCGGCAGCTGCAGCGACGACCCACGGCCTGAGAGAGC<br>CGCTGCAGCACAGGTCCCGTGCGCCCTGCCGCACAGCCTCGCCACAAC<br>CACCACcgccgccgccgccgccgccgccgccgccgccgccgccgccgccgccgccgCGCAACCC<br>GCCTCTCCCTACGGGTCCCGACTGGGCACCACTTCCGGTCCGACACGG<br>CCACGTGTTACATCTAAATGGCACCGTCCCCCGAGTGCGCCGACCTTGT<br>GCTACAGCCAGAACAGCTCCGGGAAACAAGGGCAGCGGGAGAAACGG<br>CCCGGAAAGGGACGAAGAGGCTTCCAGCTTTCGCCAAGTCGCGGCTCC |
| FRA10-499nt | TTTGAGGTCCTTCTCCCTGATGCCAAATATACCCACAGCCCTCCTCACAC<br>ACCCTTTCCCTTCCTCCCTGTGTGCCAAGTTCGCCCCCGGAGTCGTCGT<br>TTCCTTCTTTCCCGGCAGCTGCAGCGACGACCCACGGCCTGAGAGAGC<br>CGCTGCAGCACAGGTCCCGTGCGCCCTGCCGCACAGCCTCGCCACAAC<br>CACCACcgccgccgccgccgccgccgccgccgccgccgccgccgccgccgccgccgccgccgccgccg<br>ccgccgccgccgccgccgccgccgccgccgccgccgCGCAACCCGCCTCTCCCTACGGGT<br>CCCGACTGGGCACCACTTCCGGTCCGACACGGCCACGTGTTACATCTAA<br>ATGGCACCGTCCCCCGAGTGCGCCGACCTTGTGCTACAGCCAGAACAG<br>CTCCGGGAAACAAGGGCAGCGGGAGAAACGGCCCGGAAAGGGACGA<br>AGAGGCTTCCAGCTTTCGCCAAGTCGCGGCTCC |
| FRA10-550nt | TTTGAGGTCCTTCTCCCTGATGCCAAATATACCCACAGCCCTCCTCACAC<br>ACCCTTTCCCTTCCTCCCTGTGTGCCAAGTTCGCCCCCGGAGTCGTCGT<br>TTCCTTCTTTCCCGGCAGCTGCAGCGACGACCCACGGCCTGAGAGAGC<br>CGCTGCAGCACAGGTCCCGTGCGCCCTGCCGCACAGCCTCGCCACAAC<br>CACCACcgccgccgccgccgccgccgccgccgccgccgccgccgccgccgccgccgccgccgccgccg<br>ccgccgccgccgccgccgccgccgccgccgccgccgccgccgccgccgccgccgccgccgccgccgccgcc<br>gccgccgccgccgccgCGCAACCCGCCTCTCCCTACGGGTCCCGACTGGGCAC<br>CACTTCCGGTCCGACACGGCCACGTGTTACATCTAAATGGCACCGTCCC<br>CCGAGTGCGCCGACCTTGTGCTACAGCCAGAACAGCTCCGGGAAACAA<br>GGGCAGCGGGAGAAACGGCCCGGAAAGGGACGAAGAGGCTTCCAGCT<br>TTCGCCAAGTCGCGGCTCC |
| 600nt-CTrepeat | CGACGTTGTAAAACGACGGCCAGTGAATTCctctctctctctct<br>ctctctctctctctctctctctctctctctctctctctctctctctctctctctctctctctctctctctctctctctctctctctctctctc<br>tctctctctctctctctctctctctctctctctctctctctctctctctctctctctctctctctctctctctctctctctctctctctctct<br>ctctctctctctctctctctctctctctctctctctctctctctctctctctctctctctctctctctctctctctctctctctctctctctc<br>tctctctctctctctctctctctctctctctctctctctctctctctctctctctctctctctctctctctctctctctctctctctctctct<br>ctctctctctctctctctctctctctctctctctctctctctctctctctctctctctctctctctctctctctctctctctctctctctctc<br>tctctctctctctctctctctctctctctctctctctctctctctctctctctctctctctctctctctctAAGCTTGGCGTAATCATGGTCA<br>TAGCTGTT |

**Supplementary Table 1.** Detailed information for synthetic sequences.
