## Supplementary Table 2 for "Chemically Synthesized Ultra-long DNA as Building Blocks to Accelerate Complex Gene Construction in Synthetic Biology"

| Application | Primer ID | Sequences |
| --- | --- | --- |
| primers for oligo capture | APIP-Biotin | AATTATTTCTCTATCTCCCT |
| | FRA10-Biotin | TCAGGGAGAAGGACCTCAAA |
| | 600nt-Biotin | GCCGTCGTTTTACAACGTCG |
| primers for PCR after capture | APIP-F | AGGGAGATAGAGAAATAATTGCTTCT |
| | APIP-R | AAAAATACAAAATTTAGCCAGCTGTG |
| | FRA10-F | TTTGAGGTCCTTCTCCCTGATGCCAA |
| | FRA10-R | GGAGCCGCGACTTGGCGAAA |
| primers for PCR after correction | APIP-Gibson-F | acatttctctggcctaactggccAGGGAGATAGAGAAATAATTGCTTCT |
| | APIP-Gibson-R | gtctaagcttggccgccgaggccAAAAATACAAAATTTAGCCAGCTGTG |
| | FRA10-Gibson-F | acatttctctggcctaactggccTTTGAGGTCCTTCTCCCTGATGCCAA |
| | FRA10-Gibson-R | gtctaagcttggccgccgaggccGGAGCCGCGACTTGGCGAAA |
| primers for PCR of 600nt-CTrepeat | 1/2 PCR-PUC57-FP-F | GTTGTAAAACGACGGCCAGT |
| | 1/2 PCR-PUC57-FP-R | AACAGCTATGACCATGATTACGC |

**Supplementary Table 2.** Detailed information for primers.
